# Shear-induced phenotypic transformation of microglia *in vitro*

**DOI:** 10.1101/2023.02.21.529442

**Authors:** Eunyoung Park, Song Ih Ahn, Jin-Sung Park, Jennifer H. Shin

## Abstract

Brain cells are influenced by continuous fluid shear stress driven by varying hydrostatic and osmotic pressure conditions, depending on the brain’s pathophysiological conditions. While all brain cells are sensitive to the subtle changes in various physicochemical factors in the microenvironment, microglia, the resident brain immune cells, exhibit the most dramatic morphodynamic transformation. However, little is known about the phenotypic alterations in microglia in response to the changes in fluid shear stress. In this study, we first established a flow-controlled microenvironment to investigate the effects of shear flow on microglial phenotypes, including morphology, motility, and activation states. Microglia exhibited two distinct morphologies with different migratory phenotypes in a static condition: bipolar cells that oscillate along their long axis and unipolar cells that migrate persistently. When exposed to flow, a significant fraction of bipolar cells showed unstable oscillation with an increased amplitude of oscillation and a decreased frequency, which consequently led to the phenotypic transformation of oscillating cells into migrating cells. Interestingly, the level of pro-inflammatory genes increased in response to shear stress, while there were no significant changes in the level of anti-inflammatory genes. Our findings suggest that an interstitial fluid-level stimulus can cause a dramatic phenotypic shift in microglia toward pro-inflammatory states, shedding light on pathological outbreaks of severe brain diseases. Given that the fluidic environment in the brain can be locally disrupted in pathological circumstances, the mechanical stimulus by a fluid flow should also be considered a crucial element in regulating the immune activities of the microglia in brain diseases.

**Statement of Significance:** Cellular morphology and motility are important factors that encompass the alterations in protein and gene-level expressions within cells. In pathological conditions, microglia, the resident brain immune cells, are known to undergo morphodynamic transformations in response to various physicochemical stimuli. Besides the commonly known soluble biochemical factors in the microenvironment, the differential flow characteristics of ISF have been linked to several neurological diseases, such as Alzheimer’s, Parkinson’s, and brain tumors. Microglial cells, which are extremely sensitive to subtle changes in extracellular stimuli, have been identified as key players in these pathological conditions. Despite its importance, however, it has been challenging to study the sole effect of a shear flow on microglia. We investigated the morphodynamic features of microglia in response to precisely controlled interstitial-level fluid flow conditions using a microfluidic system in which isolated microglia are monitored in real-time while the undesirable effects from other extracellular factors are minimized.

## Introduction

Microglia are a type of glial cells that act as a primary immune defender in the brain. These microglia manifest high heterogeneity and plasticity in their morphological and functional features across different regions of the brain region depending on age and pathological states (1,2). Especially in response to biochemical and mechanical signals that occur in pathological conditions, microglia rapidly modify their phenotypes to initiate their neuroinflammatory or neuroprotective function (3-5). Previous studies have shown that any changes in biochemical cues such as chemokines, cytokines, adhesion molecules, and extracellular matrix (ECM) components would lead to alterations in morphology, adhesion behavior, and surface marker expressions, all of which are associated with migratory behavior of microglia (1,6-11). Moreover, microglial cells were shown to exhibit mechanosensitivity to different stiffness cues (12,13).

Recently, interstitial fluid (ISF) flow in the brain perivascular spaces, by which the parenchymal cells of the brain are surrounded, has become the subject of much interest as an essential part of brain homeostasis and pathology (**Figure 1a**) (14,15). In particular, the differential flow characteristics of ISF have been reported in many neurological disorders (16), including Alzheimer’s disease (17), Parkinson’s disease (18), and brain tumors (19). However, the microglial response to the ISF flow remains to be investigated. The complex *in vivo* brain microenvironment that includes numerous biochemical cues and intercellular interactions makes it challenging to study the sole effect of ISF on cells exclusively. Furthermore, recapitulating the extremely low flow rate of the brain in the *in vitro* experiment would require a special platform.

**Figure 1.**
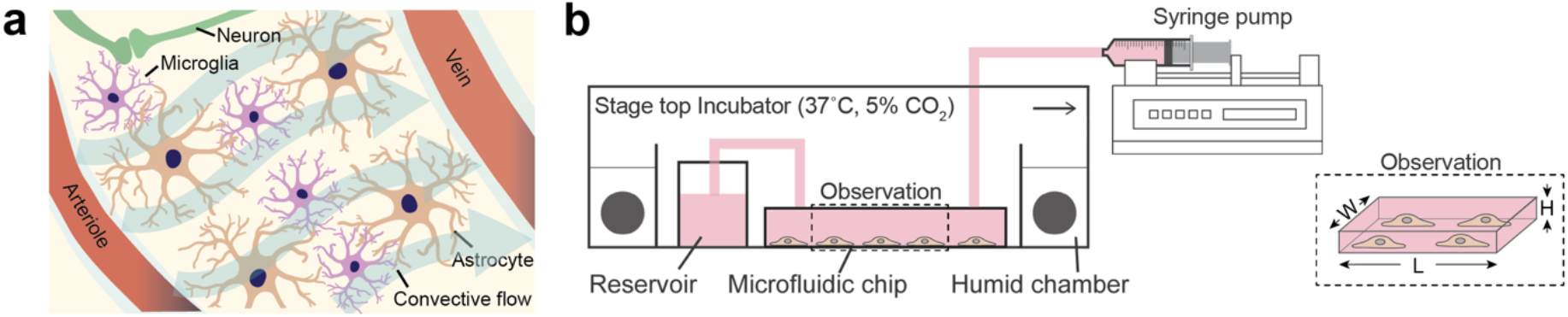
(a) Schematic showing perivascular cells under the convective bulk flow of brain interstitial fluid (ISF). (b) The schematic drawing of the microfluidic device used for our experiments. The dimension of the observation area is 3 × 0.242 × 14 mm^3^ (W × H × L). The shear flow inside a microfluidic channel was controlled by a PhD Ultra syringe pump (Harvard Apparatus, Holliston, MA, USA) in a withdrawal mode.

Here, we construct and control the brain fluidic microenvironment in microfluidic chip-based models that can reconstitute and precisely mimic the extremely low levels of ISF flow in the brain.

Using this platform, we cultured the cells at a low density of ∼ 20 cells/mm^2^ in a closed microfluidic channel system at desired flow rates to investigate the sole effect of interstitial flow. The low-density culture was to minimize undesired paracrine effects from excessively accumulated soluble factors, confirmed by the fact that the cells cultured with no flow condition did not undergo any alternations in migration modes for 6 hours of observation. In addition, under the flow conditions, the fluid flow would wash away paracrine factors, completely replacing the media with a fresh one within 10 ∼ 100 seconds, which is significantly shorter than the usual time frame of the paracrine signaling effects (∼ 5 min) (20). A fascinating phenotypic transformation was observed when the shear flow was applied to microglia cells at extremely low levels within the range of ISF (0.1 ∼ 0.3 μg min^-1^ g^-1^). In a no-flow condition, microglia exhibited two distinct morphologies with different migratory phenotypes: bipolar cells that oscillate along their long axis and unipolar cells that migrate persistently with differential centrosome location and cytoskeletal arrangement. Interestingly, introducing the shear flow induced locally oscillating microglia to transform into persistently migrating ones. Furthermore, the shear flow induced upregulation of pro-inflammatory gene expressions in microglia, implying the role of shear stress as an inherent mechanical stimulus that potentiates microglial pro-inflammatory activation in the brain. Our study demonstrates that microglia present innate heterogeneity even in the absence of extracellular stimuli, and an ISF-level stimulus can provoke a dramatic phenotypic transformation in microglia toward the pro-inflammatory states.

## Materials and methods

### Microfluidic channel fabrication and flow application

The microfluidic device used in the experiment has been described in detail previously (21). Briefly, a microfluidic device with a single microchannel (3×0.242×14 mm^3^, W×H×L) was fabricated by a conventional soft lithography process using polydimethylsiloxane (PDMS) (Sylgard 184, Dow Corning, MI, USA). PDMS mixture (10:1 elastomer base to curing agent, w/w) was degassed and poured onto SU-8 (MicroChem, MA, USA) patterned wafer and cured at 80°C for 1 h. After punching the inlet/outlet of the microchannel, the PDMS layer and a coverslip were treated with oxygen plasma for 1 min (Covance, FEMTO SCIENCE, South Korea) for irreversible bonding. The device was autoclaved for sterilization before cell culture. After 1 day of cell culture at 37°C, time-lapse imaging was turned on to capture the cells at static condition for 6 h before the steady flow was introduced. The PhD Ultra syringe pump (Harvard Apparatus, MA, USA) was set at 2, 5, and 10 *μl*⁄*min* of volume flow rate at a withdraw mode for additional 6 h before the imaging was turned off. The values of wall shear stress induced by the flow were calculated (0.002, 0.006, and 0.017 dyne/cm^2^) based on the measured maximum velocity (*u*_*max*_) of polystyrene particles in a microchannel where the focal plane was adjusted at the center of the channel height (14.449, 46.029, and 136.01 *μm*/*s*; **Fig. S1**). Here, the equation for Hagen-Poiseuille flow in a rectangular duct 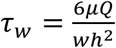 (*μ*: dynamic viscosity (7.4 × 10^−1^*kg*/(*m* ∙ *s*)), *Q*: flow rate, *w*: width of the channel (3 mm), *h*: height of the channel (0.242 mm), where *Q* = *ν*_*avg*_ *A*, and *A* = *wh*) was used to calculate the shear stress.

### Cell culture

Glial cells were isolated from the cerebral cortex of postnatal day 1-2 Sprague-Dawley rat brains (Charles River, OrientBio Inc., South Korea) following previously described methods (22,23). Briefly, after removing meninges, the cortical tissues were digested with Papain solution (LK003176, Worthington Biochem., NJ, USA) for 10 mins at 37ºC, followed by mechanical triturating to dissociate them. Then, they were passed through a 40 μm cell strainer (#93040, SPL, South Korea) and centrifuged at 1000 RPM for 5 mins. After removing the supernatant, cells were seeded in a T75 flask pre-coated with 100 μg/ml of Poly-D-Lysine hydrobromide (PDL; P6407, Sigma, MO, USA) to enhance the primary cell attachment. Next, cells were maintained in Dulbecco’s Modified Eagle’s Medium (DMEM; LM 001-05, Welgene, South Korea) supplemented with 10% heat-inactivated fetal bovine serum (FBS; S101-01, Welgene, South Korea) and 1% penicillin-streptomycin (P0781, Sigma, MO, USA).

After 7-14 days of culture, mixed glial cultures were shaken on a rotary shaker (Sejong orbital shaking incubator, South Korea) at 200 rpm for 4 h at 37°C. Primary microglia were obtained by collecting the floating cells and then cultured in the same medium used for mixed glial cell culture (22,23). Prior to seeding the cells into the microchannel, Prior to seeding the cells into the microchannel, the inner surface of the microfluidic channel was coated with 0.5 μg/cm^2^ fibronectin (#33016015, ThermoFisher, MA, USA). After 1 hour of incubation in a humidified chamber (37°C and 5% CO_2_), the channel was thoroughly washed using PBS solution three times and then by cell culture medium twice. Primary microglia were seeded at a density of 20 cells/mm^2^ to minimize direct cell-cell contact and any undesired paracrine effects from excessively accumulated soluble factors inside the channel during the observation time. The purity of the isolated microglia from mixed glia cells was > 95%, as confirmed by immunocytochemistry (**Fig. S2**).

### Time-lapse imaging and image analysis

For long-term live-cell imaging, the microchannel was placed in a microscope stage top incubator (37°C and 5% CO_2_) (**Figure. 1b**). Cell morphologies and migration were monitored in real-time for 6 h at the time interval of 1 min using an inverted microscope (Axiovert 200M, Zeiss, Germany) unless otherwise stated. To block the ROCK signaling pathway, Y-27632 (30 μM; Y0503, Sigma, MO, USA) was treated for 30 mins prior to real-time imaging.

In order to obtain the position of each cell from time-lapse images, the nuclear centroids of individual cells were tracked instead of cellular centroids to enhance the accuracy of positioning. The use of the nuclear centroid for tracking cellular motion was justified in **Fig. S3**, which demonstrates nearly identical characteristics in motion between the cellular centroid and the nuclear centroid with a similar traveled distance, amplitudes, and wavelengths but a slight phase difference of a few minutes. Cellular morphology was analyzed using ImageJ. The relative size of the two ends was obtained by dividing D_E1_ by D_E2_ for 1 hour at 2 min intervals where D_E1_ is the diameter of the circumscribed circle for a larger end and D_E2_ is the one for a smaller end at the starting point of observation (t = 0). For example, if both ends are the same size, the polarity ratio of the two ends (**R**_*E1*/*E2*_) is 1. The shape factors (S) were calculated by taking the ratio between the long axis and short axis (S = long axis/short axis) based on fit values obtained using the Fit Ellipse tool in ImageJ, and the moving directions were analyzed based on the direction of the long axis that aligns with fan-shaped lamellipodia. To analyze the intensity of the phosphorylated myosin light chain (pMLC), the cell boundary was outlined, followed by the ellipse fitting. The major axis of ellipse fitting was defined as a moving direction of a cell, and the front-rear polarity was determined by the relative size of the lamellipodium, where the front was associated with a larger lamellipodium. We then drew a line parallel to the minor axis that went through a nuclear centroid. This line serves as a reference baseline to differentiate front and rear polarity. From the fluorescence images, the fluorescence intensity of the front region was divided by the area of the front region to obtain *I*_*front*_ per unit area (*I*_*front/area*_). Similarly, *I*_*rear/area*_ was calculated by dividing the fluorescence intensity of pMLC in the rear region by the area of the rear region.

### Immunostaining

Cells were fixed with 4% paraformaldehyde (Biosesang, Korea) for 15 mins and permeabilized using 0.2% of Triton X-100 (Sigma, MO, USA) for 15 mins at room temperature. After blocking with 3% bovine serum albumin (BSA; BSAS-NZ, Bovogen, Australia) for 1 h at room temperature, the cells were incubated with primary antibodies diluted in 1 % BSA solution overnight at 4 °C: anti-rabbit IBA-1 (1:500; 019-19741, Fujifilm Wako, Japan); anti-mouse gamma tubulin (1:5000; T6557, Sigma, MO, USA); anti-mouse alpha-tubulin (1:100; A11126, ThermoFisher, MA, USA); anti-rabbit vinculin (1:50; ab129002, Abcam, UK); anti-rabbit phosphorylated myosin light chain (1:50; 3671s, Cell signaling technology, MA, USA). After extensive washing with PBS, cells were blocked with a 5 % normal goat serum (#31782, Invitrogen, MA, USA) for 30 mins at room temperature, followed by staining with secondary antibodies (Alexa Fluor 488-conjugated goat anti-rabbit IgG, 1:400, ab150077, Abcam, UK; Alexa Fluor 594-conjugated goat anti-mouse IgG, 1:400, ab150116, Abcam, UK) diluted in 1% BSA solution for 1 h at room temperature in dark. For actin staining, Alexa Fluor 568 phalloidin (1:50; A12380, Invitrogen, MA, USA) was used. DAPI was used for nuclei staining (1 µg/mL; D9542, Sigma, MO, USA) for 5 mins in dark.

### RNA extraction and real-time quantitative polymerase chain reaction (RT-qPCR)

In order to obtain a sufficient amount of mRNA for analysis, cells were cultured in a larger-scale flow-initiating device. The specific dimension of the device is described in **Fig. S4**. Device fabrication and cell culture conditions were the same as described above. mRNA of microglia was extracted using RNAiso Plus (#9109, Takara, Japan) according to the manufacturer’s instructions. Total RNA was purified using a Nanodrop spectrophotometer (ND1000, ThermoFisher, MA, USA), and cDNA was synthesized using iScript™ cDNA Synthesis Kit (#1706691, BioRad, CA, USA). RT-qPCR was performed using iQ™ SYBR green Supermix (#1708582, BioRad, CA, USA) with amplification run on a model C1000 Touch Thermal Cycler (BioRad, CA, USA). The fold change in expression was determined based on the formula 2^-ΔΔCt^, where the housekeeping gene, glyceraldehyde 3-phosphate dehydrogenase (GAPDH) was used as the standard for normalization. All primer sequences, except for Trem2 gene, are shown in the **Table S1**. Gene analysis of TREM2 (PrimePCR™ SYBR® Green Assay: TREM2 for Rat; BioRad, CA, USA) was performed following the manufacturer’s instructions.

## Results

### Two representative phenotypes of microglia in a microfluidic channel

Cells cultured in conventional culture dishes are subject to undesirable shearing events due to uncontrolled flow in large-volume Petri-type dishes (24). The shearing effect by these flows in the culture dish can be significantly greater than the ISF’s shearing effect in the brain. Thus, in the current study, we cultured microglia on a microfluidic platform that allows precise control of the flow rate while minimizing the effects of other extracellular cues (**Figure 1b**). Under static conditions in the microfluidic device, we discovered an intriguing oscillatory migratory phenotype of microglia, apart from the typical persistently migrating one (**Figure 2a, Movie S1**). While persistent microglia exhibited a run-and-turn pattern, featuring a well-developed fan-like front at its leading edge and a long tail at the trailing edge (**Figure 2b**), microglia that oscillated along their long axis had an elongated rod shape (**Figure 2c**). During this oscillation, the active ruffling in the lamellipodium at both ends was observed at a length scale comparable to that of a cell body. In addition to bipolar oscillating and unipolar migrating cells observed in a microchannel, the third type of multipolar cells was observed when the cells were cultured in a conventional TCPS dish (**Movie S2 and Fig. S5**). Since these multipolar microglia characterized by a repeated extension and retraction of lamellipodium at multiple sites were not found in the microchannel setup with a controlled flow condition, we focused on two phenotypes, namely oscillating and migrating, by following the trajectories of their nuclei over a one hour. Unlike the steady translocation of nuclei in persistently migrating cells (**Figure 2d**), oscillating cells’ nuclei moved in a sinusoidal fashion relative to one central point (*y* = 0), whose value was defined by the mean of the nuclear position during the observation time (**Figure 2e**).

**Figure 2.**
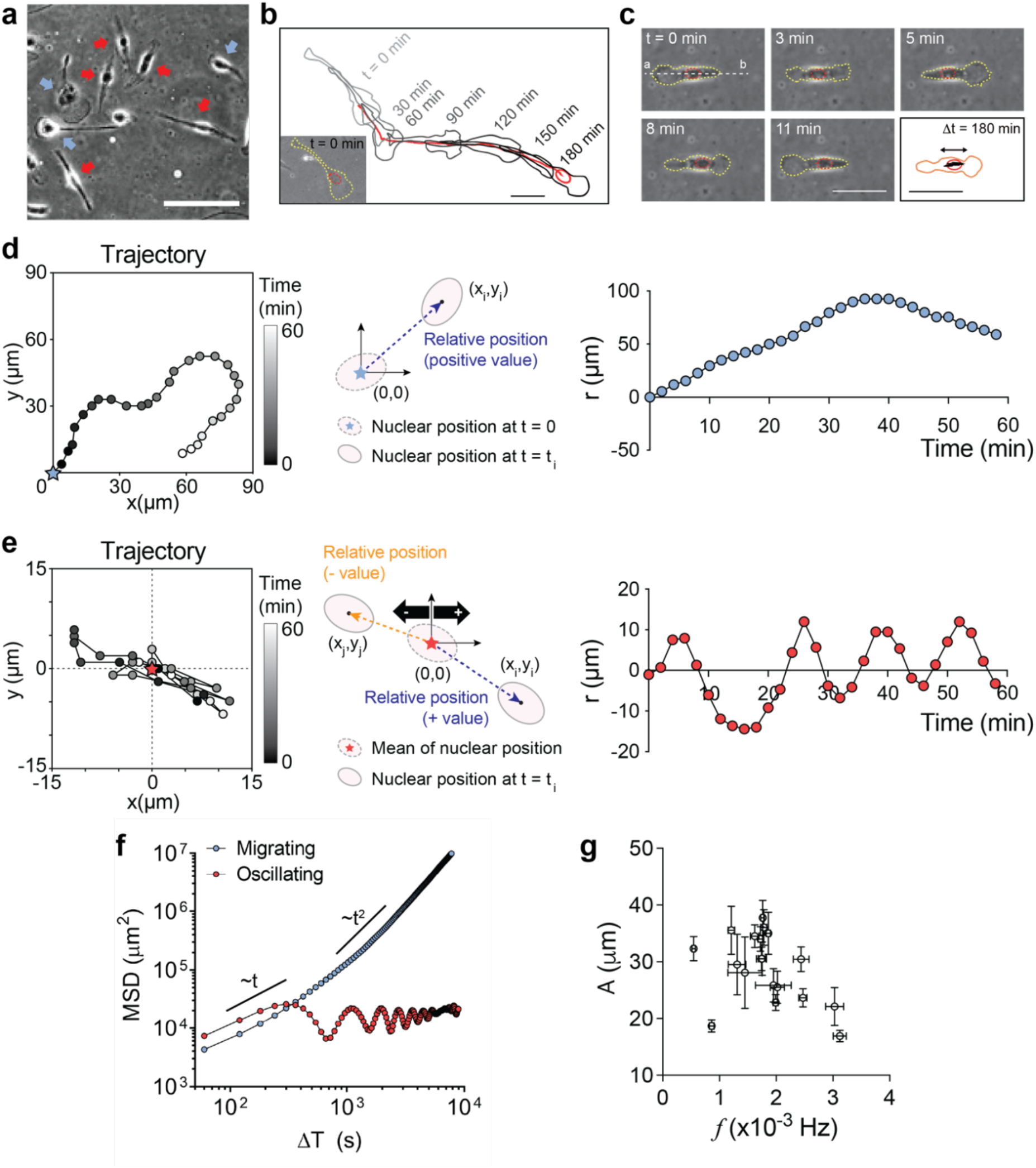
Two representative phenotypes of microglia were observed in a microfluidic channel without a shear flow. (a) Snapshot image of the representative migrating (blue arrow) and oscillating microglia (red arrow). Scale bar: 50 μm. (b) Moving trajectory of migrating microglia showing a persistent migration. The cell outline at each time point is overlaid for visual guidance. Scale bar: 50 μm. (c) Sequential images of the representative oscillating microglia showing an oscillating motion from t = 0 to 11 mins and its trajectory obtained by tracking the center location of the cell nucleus for 180 mins (black line in right-bottom image). Here, the magnitude of the arrow indicates the oscillating amplitude of the cell. Scale bar: 50 μm. (d, e) The typical trajectory of migrating microglia (d) and oscillating microglia (time interval: 2 mins) (e). (f) Double log plots of MSD vs. Δ t of the two-representative migrating (blue) and oscillating microglia (red). (g) Graph of oscillating amplitude (*A*) vs. frequency (*f*) of oscillating microglia (n = 18, * *p* < 0.05 by student t-test). Each error bar is the standard deviation for each data set.

Their mean-squared displacements (MSD) profiles revealed their characteristic motion with migrating microglia exhibiting persistent migration with its MSD curve in a slope (α) between 1 and 2, whereas oscillating microglia exhibited oscillation with limited motile displacement following a brief random migration (α ≈ 1) (Figure 2f). Despite their short migration distance, oscillating cells moved significantly faster than migrating cells (Figure 1g), with their stable yet dynamic oscillation. In particular, the oscillating motion was maintained stably over the period of six hours at a frequency of 0.003 ± 0.0001 Hz and an amplitude of 9.0 ± 0.9 μm (**Figure 2g**). Moreover, the amplitudes of the oscillating cells (A) were inversely proportional to the oscillation frequencies (f) (**Figure 2g**).

### Migratory phenotypes of microglia represented by their morphological features

Cell migration is accompanied by morphological changes in cells. In general, migrating cells exhibit lamellipodium on the front side of their direction of movement. To establish a distinction between the two migratory phenotypes based on their morphology, we analyzed the lamellipodial dynamics of the cells. To quantify the sizes of both ends of the cell body, two individual-circumscribed circles were drawn on both leading and trailing sides, and then the polarity ratio of the two diameters of the circumscribed circle was calculated (**Figure 3a**). Here, the circumcircle with a larger diameter was designated End1 (***D***_*E*1_), and one with a smaller diameter was designated End2 (***D***_*E*2_) based on the data at t = 0. For example, if both ends are the same size, the polarity ratio of the two ends (**R**_*E*1/*E*2_) would be 1. Migrating cells with a stable polarity maintained a larger circumcircle at one end, as indicated by their **R**_*E*1/*E*2_ value being greater than 1 (**Figure 3b**). On the other hand, the **R**_*E*1/*E*2_ of oscillating cells oscillate symmetrically across the baseline where **R**_*E*1/*E*2_ equals 1 (log_/5_ **R**_*E*1/*E*2_ = 0) (**Figure 3c**). Interestingly, the period of the **R**_*E*1/*E*2_ oscillation was found to be linearly correlated to the period of nuclear oscillation (α = 0.6545, R^2^ = 0.844) (**Figure 3d** and **Fig. S6**).

**Figure 3.**
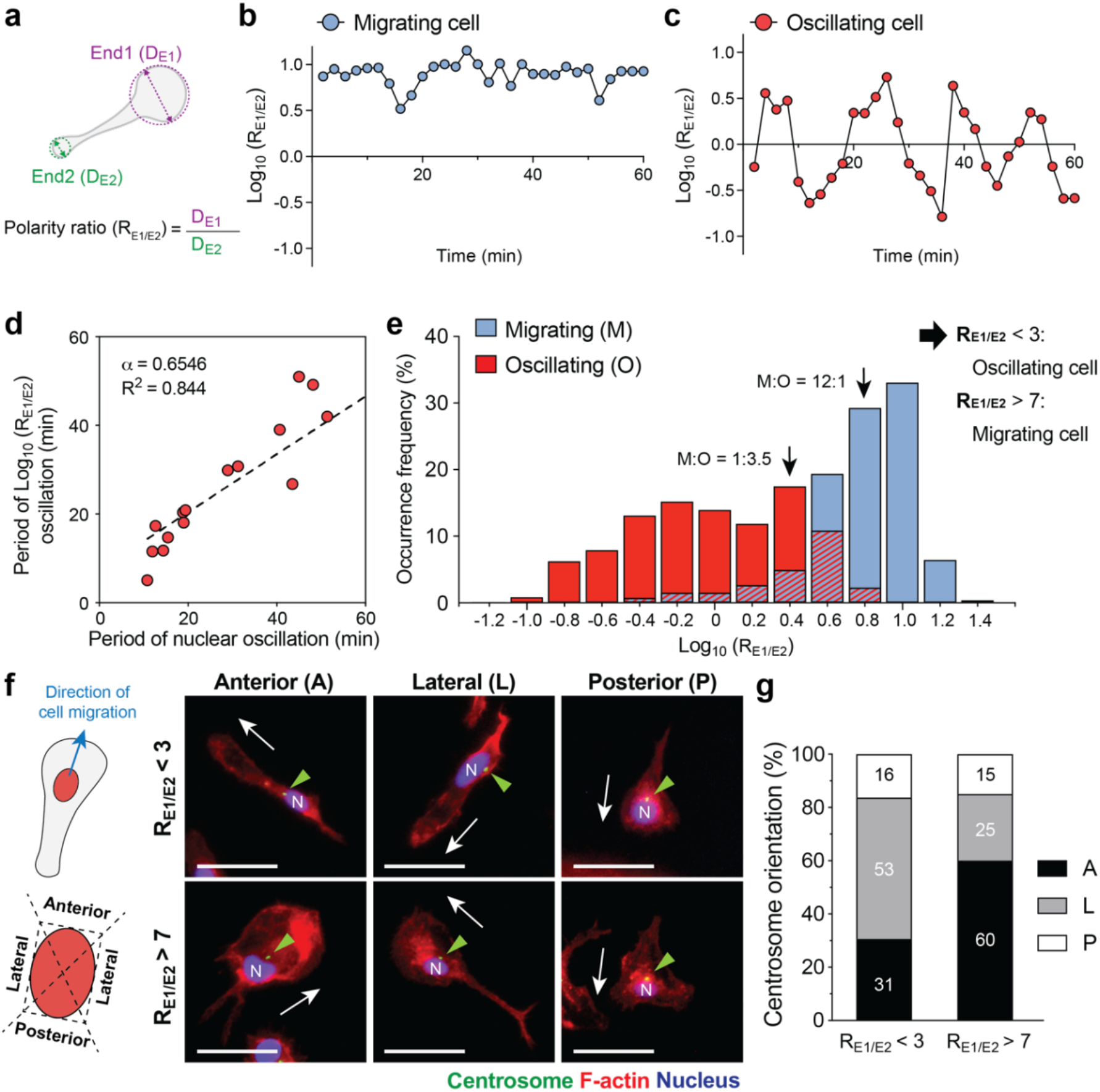
Discrimination of two migratory behaviors based on the morphology and the centrosome position. (a) Schematic description of circumcircles drawn at both end sides of a cell boundary (*E*1 and *E*2). (b,c) The ratio between the diameters of two circumcircles (**R**_*E*1/*E*2_) measured at both sides of migrating (b) and oscillating microglia (c) (time interval: 2 mins). (d) The oscillating period in nuclear translocation (**Figure 2e**) versus the **R**_*E*1/*E*2_ graphs (**Figure 3c**) obtained from oscillating cells. (*See* ***Fig. S6*** *for details*) (e) Occurrence frequencies showing the distribution of the polarity ratio (**R**_*E*1/*E*2_) measured for migrating (blue) and oscillating microglia (red) for 1 hour at 2 min intervals. The left arrow in the graph indicates the critical polarity ratio below which the oscillating cells are predominant (the probability of being an oscillating cell is more than 78%). Similarly, the right arrow indicates the critical polarity ratio over which the migrating cells are predominant (the probability of being a migrating cell is more than 98%) (n = 16 and 9 for oscillating and migrating cells, respectively). (f) Representative immunostaining images showing the locations of centrosomes in microglia (F-actin, red; γ-tubulin, green (indicated by arrowheads); DAPI, blue). White arrows indicate the expected moving direction of microglia based on the position of fan-shaped lamellipodia. Scale bar: 50 μm. (g) The percentage of microglia having centrosomes placed in anterior (A), lateral (L), and posterior (P). The quantification was performed based on the immunostaining images of F-actin, γ-tubulin, and DAPI (n = 20 and 98 for cells with **R**_*E*1/*E*2_ > 7 and **R**_*E*1/*E*2_ < 3, respectively).

The threshold criterion was then determined using occurrence frequencies of **R**_*E*1/*E*2_ obtained from all cells over one hour in order to quantitatively discriminate between the two migratory phenotypes (**Figure 3e**). Oscillating cells showed symmetrically distributed occurrence frequencies with its center near 0 (**R**_*E*1/*E*2_ ∼1), indicating that cells with similar end sizes were frequently observed (**Figure 3c**). On the other hand, the occurrence frequencies of **R**_*E*1/*E*2_ in migrating cells showed a peak at 1 (**R**_*E*1/*E*2_ = 10), indicating that migrating cells had ∼10-fold larger end than the opposite side (**Figure 3b**). Based on these differential distributions of **R**_*E*1/*E*2_ frequencies, we assumed that the cells of **R**_*E*1/*E*2_ < 3 were oscillating cells with >78% certainty. Similarly, cells with **R**_*E*1/*E*2_ > 7 were classified as migrating cells with > 92% certainty.

The centrosome, which is typically located behind the leading edge of a migratory cell, regulates directional cell migration (25). In particular, the fan-shaped lamellipodia represents the leading edge of a migrating cell; thus, the instantaneous direction of cell migration in the immunofluorescent images was accordingly determined. Therefore, we classified the position of the centrosome (gamma-tubulin; γ-tubulin) into anterior, lateral, and posterior with respect to the nuclear position and the direction of cell migration (**Figure 3f**). Over 60% of the migrating cells (**R**_*E*1/*E*2_ > 7) featured anteriorly oriented centrosomes, whereas over 50% of the oscillating cells (**R**_*E*1/*E*2_ < 3) exhibited laterally oriented centrosomes (**Figure 3g**). These findings suggest that the dynamic migratory characteristics of microglia can be interpreted based on the static morphological features, including the ratio of the cell body’s both ends and the position of the centrosome.

### Distinct myosin distribution in the two phenotypes of microglia

Cell migration generally consists of four sequential steps: lamellipodial protrusion at the leading edge, cell adhesion, rear-end contraction, and detachment of focal adhesions at the rear end (26). More importantly, these cell migration steps are regulated by cytoskeletal dynamics, which include actin and microtubule remodeling, focal adhesion turnover, and myosin activity (27). Thus, we compared the cytoskeletal arrangements of oscillating and migrating microglia in order to identify the critical cytoskeletal components involved in the two distinct microglial migratory phenotypes, defined by the polarity ratio of both ends (**R**_*E*1/*E*2_) (**Figure 3e**). Both oscillating and migrating cells exhibited cortical actin structures at their periphery and lacked actin stress fibers, implying that microglia do not migrate in the conventional mesenchymal mode (**Figure 4a**) (28). Similarly, microtubules, a critical molecule that regulates the direction of cell migration (25), were not organized regardless of the migratory phenotypes, implying that migration of both phenotypes is not controlled by the polarity of the microtubule network (**Figure 4a**). Moreover, both oscillating and migrating microglia did not show apparent vinculin dots at the cell periphery, suggesting that vinculin was not likely to be a key adhesion molecule to differentiate the two modes of migration. (**Figure 4b**).

**Figure 4.**
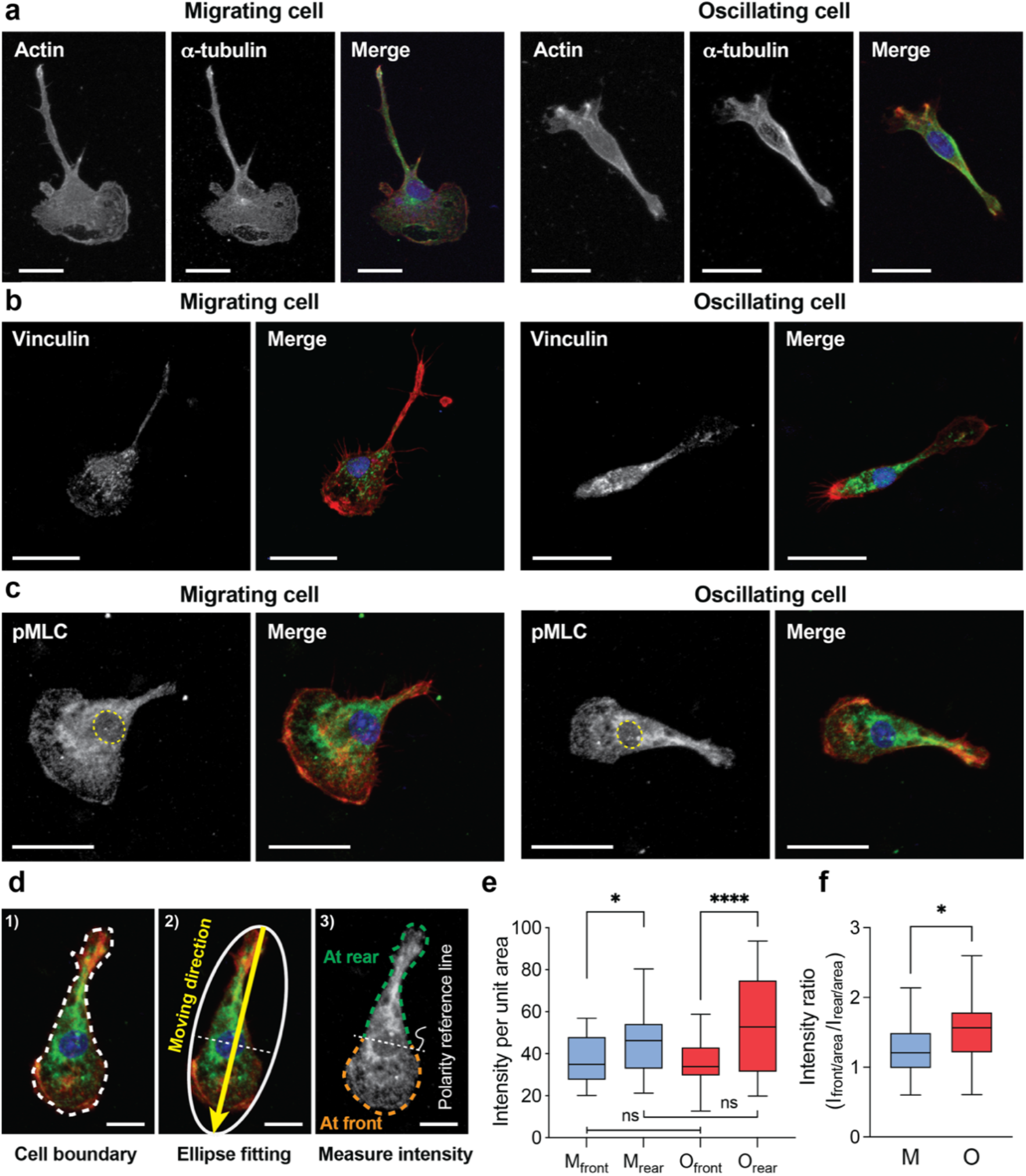
Distribution of cytoskeleton and associated proteins in migratory phenotypes of microglia. (a-c) Representative immunostaining images showing microtubule (alpha-tubulin; green) (a), vinculin (green) (b), and phosphorylated myosin light chain (pMLC; green) (c) in fixed microglia. Cells were co-stained with F-actin (phalloidin; red) and DAPI (blue) (scale bars = 25 µm). (d) Quantitative analysis of the pMLC distribution within a cell: (1) delineate cell boundary, (2) define the moving direction by ellipse fitting, (3) divide the cell into front and rear with respect to the centroid of the nucleus and measure the sum of the fluorescence intensity in front and rear side, respectively (scale bar = 10 µm). (e) The intensity per unit area at the front (*I*_*front/area*_) and rear (*I*_*rear/area*_) region of oscillating and migrating cells. (f) The pMLC intensity ratio (*I*_*rear/area*_ / *I*_*front/area*_) in oscillating and migrating cells. (n = 17 and 18 for oscillating and migrating cells in **Figures e** and **f**; *p<0.05 and ****p<0.0001 by student t-test).

Thus, we hypothesized that microglial motility is achieved by the cyclic expansion and contraction of the cell body, as indicated by the intracellular distribution of phosphorylated myosin light chain (pMLC), a key actuator of myosin II. We investigated the intracellular distribution of pMLC in relation to the position of the nucleus. In both migrating and oscillating cells, pMLC was predominantly distributed around the nucleus (**Figure 4c**). Now the fluorescence intensity of the front region was divided by the area of the front region to obtain *I*_*front*_ per unit area (*I*_*front/area*_). Similarly, *I*_*rear/area*_ was calculated by dividing the fluorescence intensity of pMLC in the rear region by the area of the rear region (**Figure 4d**). Here, the front-rear polarity was determined by the relative size of the lamellipodium, where the front was associated with a larger lamellipodium. As shown in **Figure 4e**, the pMLC intensity per unit area was significantly higher on the rear side than the front side, regardless of the migration modes. Interestingly, the relative pMLC intensity in the rear compared to that of the front side was significantly more significant for the oscillating cells (**Figure 4f**), suggesting the possible involvement of higher pMLC-mediated contractility in actively oscillating cells. However, one caveat of this analysis is that the front vs. rear polarity determination from the fixed cells could inherently have some errors, requiring live-imaging-based analysis for better accuracy.

### Shear-induced phenotypic transformation of microglia

Microglia constantly monitor their immediate local environment and respond rapidly to any disturbance in the microenvironment’s homeostasis. One significant yet understudied source of disturbance in the brain is the ISF produced by the cerebrospinal fluid (CSF) and the vascular system. ISF causes shear stress on microglia, and the degree of shear stress varies according to the degree of glymphatic dysfunction present in brain pathology (16,29). Using our microfluidic device that enables precise flow control, we monitored microglia’s morphological and migratory phenotypes in real time for six hours under different levels of shear stress. We discovered that when shear stress was applied at 0.006 and 0.017 dyne/cm^2^, the ratio of oscillating to migrating microglia decreased, whereas the number of oscillating cells was threefold of migrating cells in a static condition (**Figure 5a**). The transition from oscillating to migrating microglia began to occur within an hour after shear stimulation and continued to occur over 6 hours of observation (**Fig. S7)**. Not only did shear stimulation (> 0.017 dyne/cm^2^) increase the relative number of migrating microglia, but it also increased persistence and migration speed (**Figure 5b-c**). However, the increase in the number of cells with anterior centrosomes was not as significant as the increased population of migratory cells under flow, implying that the centrosome location may not be a direct indicator of transformed migratory cells from the oscillating cells. Instead, the centrosome repositioning must occur after the transformation of oscillating cells into migratory cells (**Fig. S8**). However, the cells did not display any biased directedness of cell migration in response to the flow direction (**Figure 5d**).

**Figure 5.**
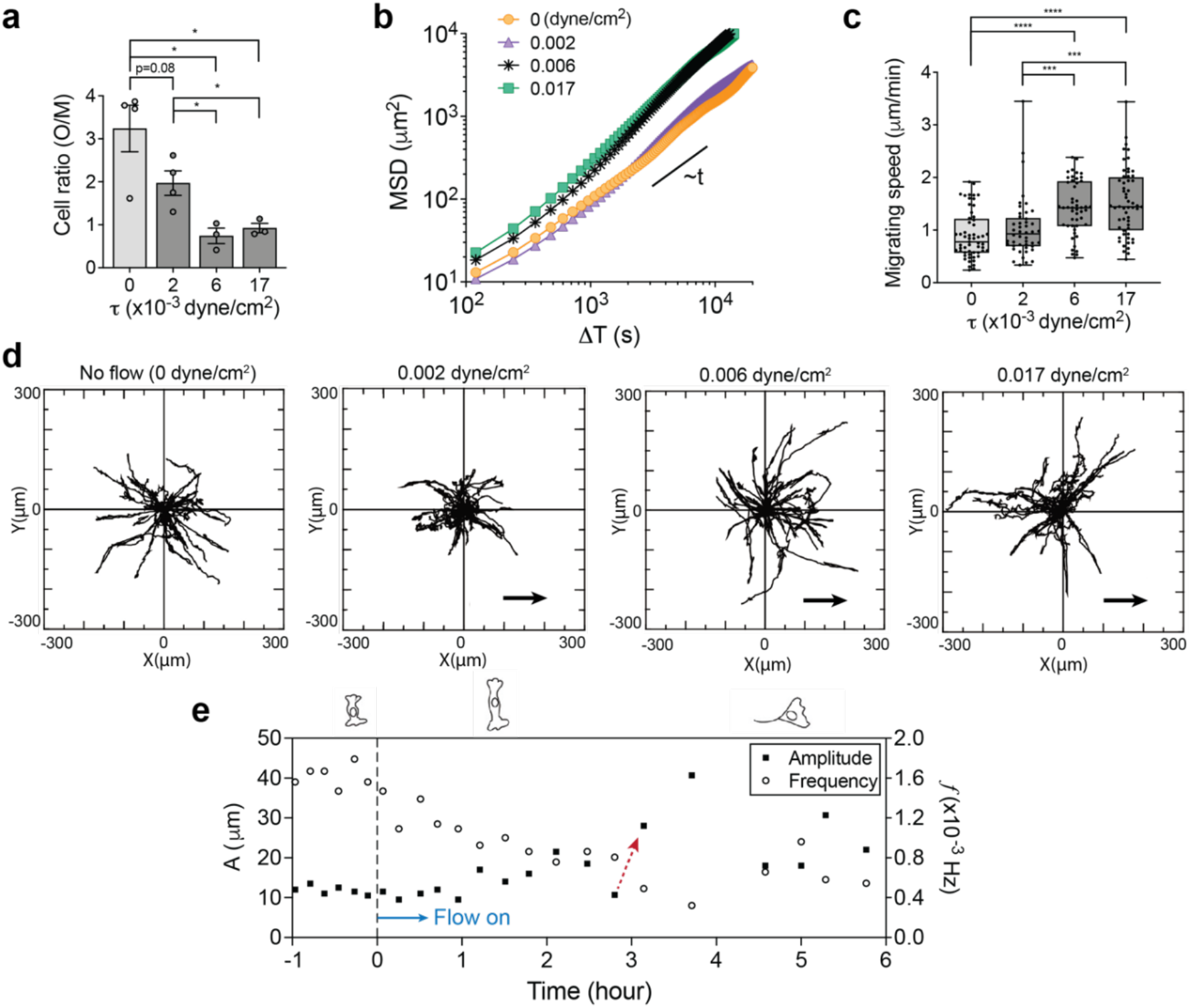
The shear-induced transformation from oscillating to migrating microglia. (a) Changes in the ratio of oscillating (O) to migrating cells (M) with an exposure to shear stress over 6 h, showing significantly reduced O/M ratio from 2.97 ± 0.39 (τ = 0) to 0.92 ± 0.08 (τ = 0.017 dyne/cm^2^) (n = 4 for 0 and 0.002 dyne/cm^2^, n=3 for 0.006 and 0.017 dyne/cm^2^, * *p* < 0.05 by student t-test, Error bar: standard error for each experiment). (b) Double log plot of MSD vs. ΔT of single migrating microglia observed in no flow and flow conditions. (c) Shear stress-dependent migrating speed (n=57 for 0, n= 46 for 0.002, n=49 for 0.006, n=56 for 0.017 dyne/cm^2^, *** *p* < 0.01 and ****p<0.001 by student t-test). (d) The trajectory patterns of a single migrating microglia were observed in no flow and flow conditions, showing no directional preference upon shear stimulation. Here, the flow direction was indicated by arrows. (e) The percentage of microglia having centrosomes in anterior, lateral, and posterior in the absence and presence of shear flow analyzed from the immunostaining images of F-actin, γ-tubulin, and DAPI. (n = 165 and 212 for τ= 0 and 0.017 dyne/cm^2^, respectively) (f) Changes in the oscillating amplitude (*A*) and frequency (*f*) of oscillating microglia observed during its transitional period, indicating a transformation at ∼ 4 h (red dotted arrow).

Furthermore, in response to shear flow, oscillating microglia exhibited a gradual increase in oscillation amplitude as frequency decreased, indicating an unstable oscillation followed by a transformation into the migrating phenotype (**Figure 5f** and **Movie S3**). This phenotypic transformation of microglia was also demonstrated by its morphological change, as defined by the shape factor (S = ratio of the long axis/short axis), which was gradually increased over time until one end of the cell body detached from the surface to initiate persistent migration (**Fig. S7**). Taken together, shear stress transformed oscillating cells into persistent migrating cells by actuating them to escape from the original oscillating region defined by the cell body length. The potential effects of shear stress on pMLC redistribution of microglia were demonstrated by the relative change in the ratio of oscillating to migrating cells following inhibition of the cell’s Rho-associated protein kinase (ROCK)-signaling pathway (**Additional files: Fig. S8 and Movie S4**).

As represented by pro- and anti-inflammatory paradigms, inflammatory states of microglia are intimately linked to the changes in the morphological and migratory states of the cells (30). We analyzed the expressions of inflammatory genes in microglia with or without flow to examine the immediate effect of shear stress on the cells’ inflammatory responses. The shear-induced transition began to occur 1 hour after applying the shear stimuli and continued to occur for the rest of the observation time (**Fig. S7**). We chose 6 hours after the onset of the shear flow for the PCR time point to be inclusive of all events that have occurred during the observation time. At this time point, more than 50% of the cells exhibited the migratory phenotype (**Figure 5a**). After six hours of shear stress at 0.017 dyne/cm^2^, cells expressed increased levels of pro-inflammatory genes, interleukin 1β (IL-1β), and tumor necrosis factor α (TNF-α) (**Figure 6a)**.

**Figure 6.**
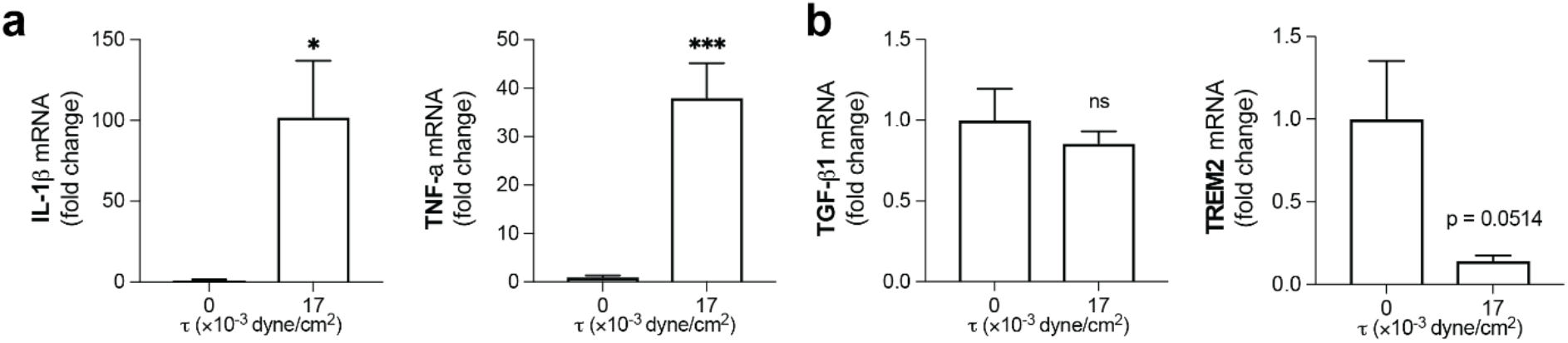
Inflammatory gene expression after 6 h of culture under static conditions and shear stress. In response to τ=0.017 dyne/cm^2^ of shear stress, microglia showed increased levels of pro-inflammatory genes (Il-1β and TNF-α) (a) and no significant changes in anti-inflammatory genes (TGF-β1 and Trem2) (b) (n > 4, *p < 0.05 and *** p < 0.001 by student t-test). For raw data, see **Table S2**.

By contrast, the expression levels of anti-inflammatory genes, represented by transforming growth factor-beta1 (TGF-β1) and triggering receptor expressed on myeloid cells 2 (TREM2), exhibited only little change (**Figure 6b**). Our findings suggest the shear flow-induced activation of microglia towards pro-inflammatory states. Although the qPCR analysis with a few selected gene sets may be insufficient, our results demonstrated the potential effects of the disrupted brain ISF on the alternations in microglial reactivity during pathophysiological responses in the brain.

## Discussion

In the brain perivascular spaces, brain parenchymal cells, including neurons, microglia, and astrocytes, are surrounded by ISF (**Figure 1a**). The bulk flow of ISF is generated primarily by fluids from the brain vasculature, inducing shear stress on the brain parenchymal cells. In this study, we investigated phenotypic heterogeneity and plasticity of microglia in response to fluidic shear stress in a microfluidic channel, where the cells can be monitored in real-time in a precisely controlled microenvironment. Our microfluidic device, in conjunction with a custom-built microscope stage top incubator and a programmable syringe pump, enabled real-time monitoring of cells over a six-hour period, and the actual flow rate applied in the microchannel was determined by tracking the movement of microbeads. Additionally, the morphologies, migrating patterns, centrosome location, and cytoskeletal structure of microglia were used to characterize the distinct phenotypes of microglia (**Table 1**).

**Table 1.** Distinct phenotypes of oscillating and migrating microglia.

|  | Oscillating microglia | Migrating microglia |
| --- | --- | --- |
| **MSD profile (MSD vs. lag time)** | Linear slope between 1 and 2 | Linear slope around 1 with oscillating tails |
| **$R_{E1/E2}$** | Smaller than 3 | Larger than 7 |
| **Centrosome position** | Laterally oriented in over 50% | Anteriorly oriented in over 60% |
| **Ratio of fluorescence intensity**<br>($I_{rear/area} / I_{front/area}$) | Significantly higher in oscillating cells than that in migrating cells | |

Because microglia cells in the brain are exposed to various extracellular stimuli, it is difficult to investigate the effects of the individual stimulus separately. Moreover, the unique sensitivity of microglia requires a precisely controlled, flow-free platform for studying their phenotypes in the absence of external cues by the flow. In a static condition established in our microfluidic device, we classified the microglia into two distinct modes based on their motile characteristics; oscillating and migrating microglia. We further demonstrated that their morphology-based phenotypic differences could serve as a quantitative criterion for distinction. Moreover, we confirmed that these two different phenotypes of microglia exhibited remarkable differences in the intracellular distribution of pMLC.

ISF in the perivascular spaces of the brain is extremely slow with a flow rate in a range of 0.1 ∼ 0.3 μg min^-1^ g^-1^ (31). Our microfluidic platform enabled us to generate and control the slow flows that mimic the ISF. Even at a low flow rate, microglia exhibited drastic changes in response to shear stress, transitioning from oscillating to migrating phenotypes with a much greater sensitivity to the shear flow compared to other cell types (21,32). Moreover, shear stress induced increased motility with increased persistence and cell migration speed without any directional preference. In addition to the morphological and motility changes, we confirmed the enhanced expressions of two pro-inflammatory marker genes (TNF-α and IL-1β) in microglia, supporting the shear flow-induced activation of microglia towards pro-inflammatory states. However, the mechanism underlying the pro-inflammatory activation of microglia in response to shear stress remains elusive. Furthermore, besides various soluble biochemical modulators known to regulate the signaling pathways in the phenotypic transition of microglia (33), our study suggests that the brain ISF can be a potential upstream stimulator for phenotypic transition in microglia, playing a critical role in immune responses during pathophysiological conditions in the brain.

Taken together, our findings indicate that the shear stress induced a phenotypic transition in microglia, possibly toward pro-inflammatory states, playing critical roles in brain pathology. We believe our study may shed new light on the potential effects of the disrupted brain ISF on the microglial reactivity during pathophysiological conditions in the brain.

## Appendices

Data are presented as the mean ± the standard error of the mean (SEM) unless otherwise stated. All statistical analyses were done using GraphPad Prism 6 (GraphPad Software Inc., La Jolla, CA, USA). Unpaired Student’s t-tests were applied, and p-values of < 0.05, 0.01, 0.005, and 0.001 were considered significant in all tests.

## Author Contributions

Designed research: S. I. Ahn, E. Park, J.-S. Park, J. H. Shin. Performed research: E. Park, S. I. Ahn, J.-S. Park. Contributed analytic tools: S. I. Ahn, E. Park. Analyzed data: S. I. Ahn, E. Park, J.-S. Park, J. H. Shin. Wrote the paper: S. I. Ahn, E. Park, J.-S. Park, J. H. Shin.

## ACKNOWLEDGMENTS

This work was supported by the National Research Foundation of Korea (NRF) of the Ministry of Science and ICT (NRF-2021R1A4A1031198). We thank Prof. K. J. Lee for his support in cell culture.

