## Supplemental Files for "Shear-induced phenotypic transformation of microglia *in vitro*"

#### CONTENTS

**Figure S1**. The graph of the flow rate of the syringe pump (Q) vs. the average velocity ($u_{max}$) of polystyrene (PS) particles (600 nm diameter) in a microchannel

**Figure S2**. Purity of isolated primary microglia.

**Figure S3.** The characteristics of cell nuclei and the centroid of cell bodies of migrating and oscillating microglia.

**Figure S4.** Details of the scaled-up channel used for analyzing the gene expressions after shear stimulation.

**Figure S5.** Three representative phenotypes of microglia observed when cultured in a tissue culture polystyrene (TCPS) dish.

**Figure S6.** Dynamic features observed in the nucleus (a) and both end circumstance (b) in the oscillating microglia, corresponding to those of graphs in Figures 2e and 3c, respectively.

**Figure S7.** The transition from oscillatory to migratory microglia under the shear flow applied at 0.017 dyne/cm$^2$.

**Figure S8.** The orientation frequencies of the centrosome in microglia in the absence and presence of shear flow were analyzed from the immunostaining images of F-actin, g-tubulin, and DAPI.

**Table S1.** Primer sequences for gene analysis.

**Table S2.** Fold change calculation.

**Movie S1**. Typical motions of microglia in a no flow condition. Time-lapse live-cell images were taken for 420 mins. Scale bar: 100 μm.

**Movie S2**. Motions of microglia when cultured in a tissue culture polystyrene (TCPS) dish. Time-lapse live-cell images were taken for 100 mins. Scale bar: 100 μm.

**Movie S3.** Typical transformation of oscillating to migrating cells under fluid flow at $\tau$ = 0.002 dyne/cm$^2$. Flow was turned on at t = 0 min. Time-lapse live-cell images were taken for 6h. Scale bar: 100 μm.

#### FIGURE S1

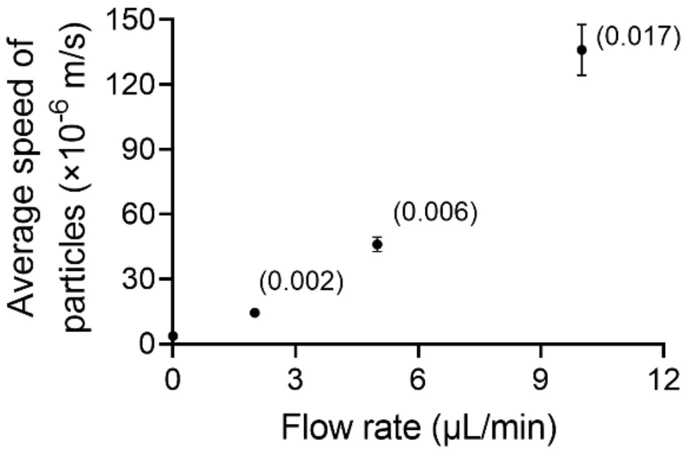


| $\mathbf{Q}\ (\boldsymbol{\mu l/min})$ | 0 | 2 | 5 | 10 |
|---|---|---|---|---|
| $\mathbf{Q}\ (\times \mathbf{10^{-11}}\ \boldsymbol{m^3/s})$ | 0 | 3.333 | 8.334 | 16.667 |
| $\boldsymbol{u_{max}}\ (\times \mathbf{10^{-6}}\ \boldsymbol{m/s})$ | $3.7078 \pm 3.1943$ | $14.449 \pm 4.5185$ | $46.029 \pm 10.206$ | $136.01 \pm 33.324$ |
| **N** | 3 | 6 | 9 | 8 |

**Figure S1.** The graph of the flow rate of the syringe pump (Q) vs. the average velocity ($u_{max}$) of polystyrene (PS) particles (600 nm diameter) in a microchannel where the focal plane was adjusted at the center of the channel height. Here, the magnitudes of the shear stress (τ) for each flow rate are written in brackets (unit: dyne/$cm^2$). (Error bars: standard error means for each data set)

### FIGURE S2

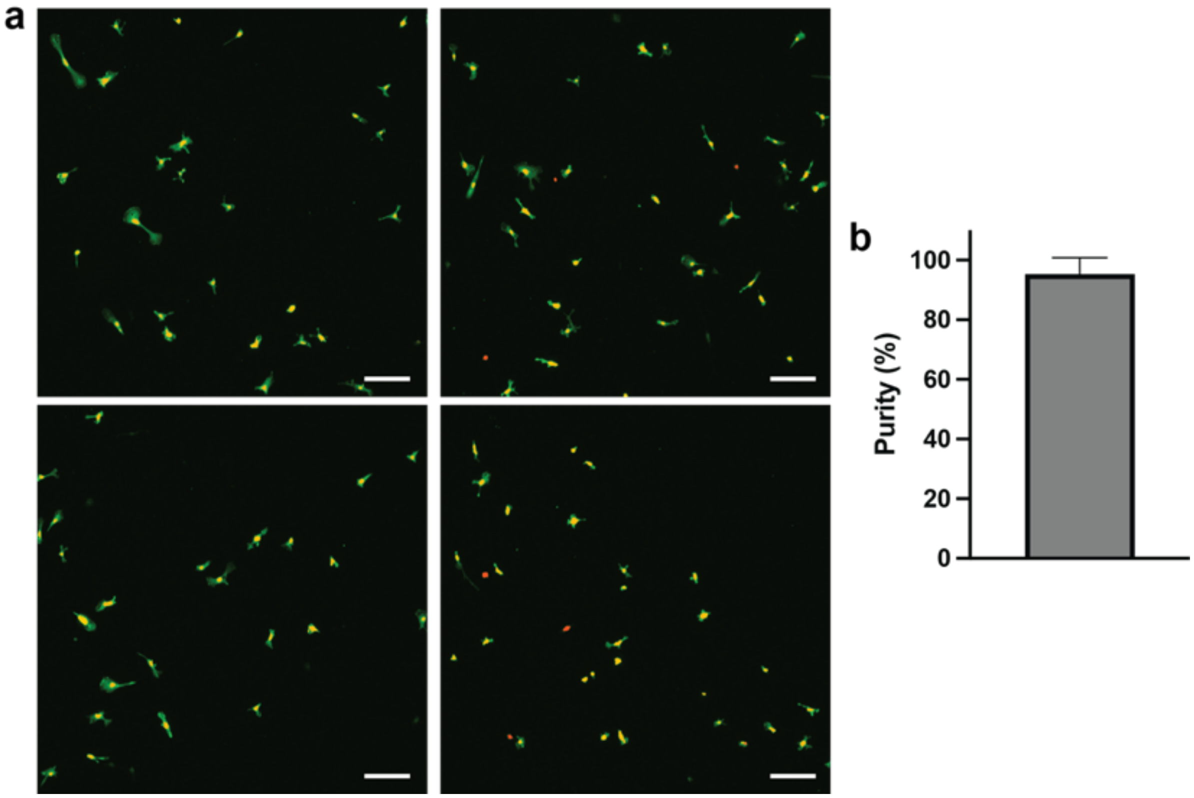


**Figure S2.** Purity of isolated primary microglia. (a) Immunostaining of isolated microglia (IBA-1; green) and nuclei (DAPI; red). Scale bar = 100µm. (b) 93 out of total 97 cells were IBA-1 positive, so the culture purity was 95.2 ± 2.7 %, calculated as IBA-1$^{+}$/DAPI$^{+}$ (%).

#### FIGURE S3

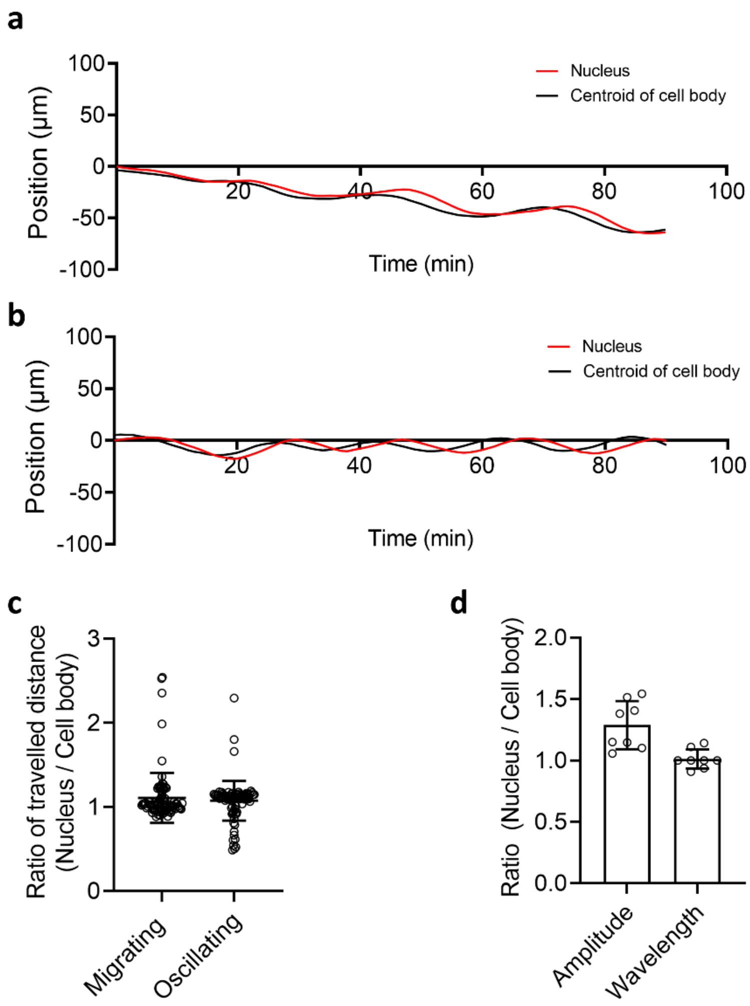


**Figure S3.** (a, b) The positions of cell nuclei and the centroid of cell bodies of migrating (a) and oscillating microglia (b). (c) The ratio of the traveled distance of cell nuclei to the centroid of cell bodies. (d) The amplitude and wavelength ratio of a nucleus to the centroid of the cell body of an oscillating cell.

### FIGURE S4

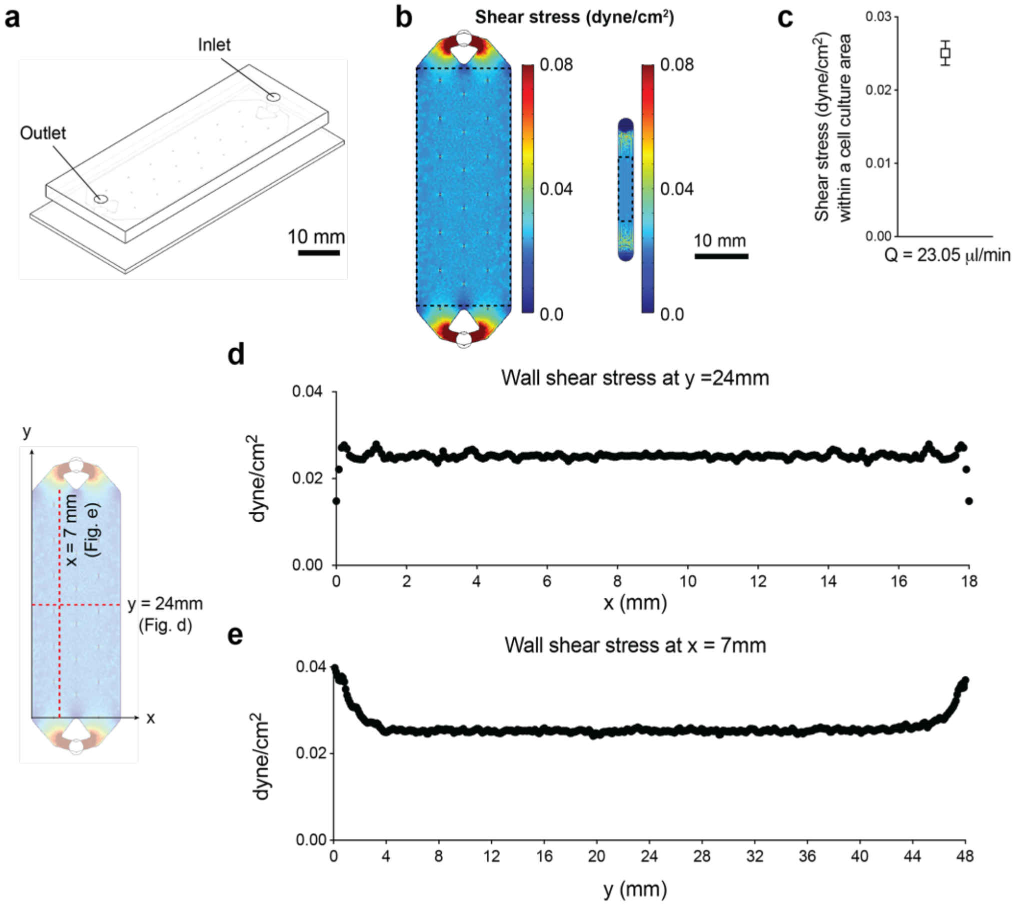


**Figure S4.** Details of the scaled-up channel used for analyzing the gene expressions after shear stimulation. (a) The schematic of a scaled-up channel. This channel has one inlet and one outlet. The projected area of this channel is 20 times larger than the microfluidic chip that has been used for live-imaging analysis in this study (**Figure 1b**). ($18\times0.15$ mm$^2$ (W×H)) (b) Simulation results of the fluid shear stress in a scaled-up channel. In this simulation, under the same flow rate to each channel, we confirmed that the level of fluid shear stress in the scaled-up channel follows the equation in the previous study (1). In addition, a scaled-up microchannel (left) was shown to generate a uniform level of fluid shear stress to the cell culture area (black dotted box), similar to the original microfluidic chip (right, black dotted area). (c) The average shear stress within the observation area in accordance with the introduction of the flow rate ($0.025 \pm 0.0016$ dyne/cm$^2$). (d-e) The velocity profiles in x and y directions. (d) in y = 24mm (d) and x = 7 mm (e) positions.

### FIGURE S5

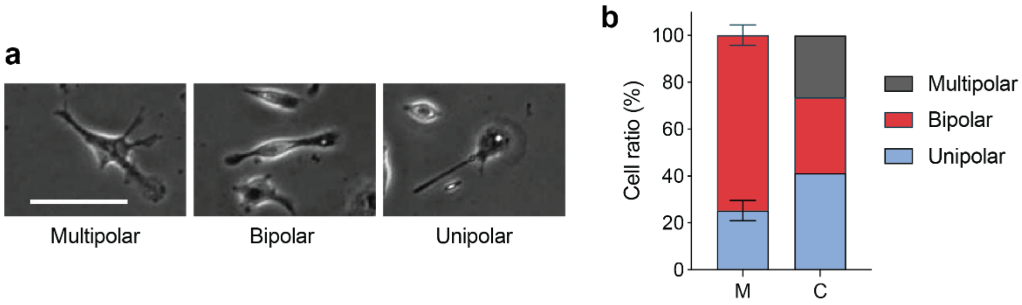


**Figure S5.** Three representative phenotypes of microglia observed when cultured in a tissue culture polystyrene (TCPS) dish (SPL, #20035). (a) Besides two typical phenotypes of oscillatory and migratory microglia observed in a microfluidic channel, multipolar microglia with an irregular shape co-exist as a third fraction. Scale bar = 100 μm. (b) The ratio of three and two different phenotypes of microglia estimated in a microfluidic channel (M) and a conventional culture dish (C).

#### FIGURE S6

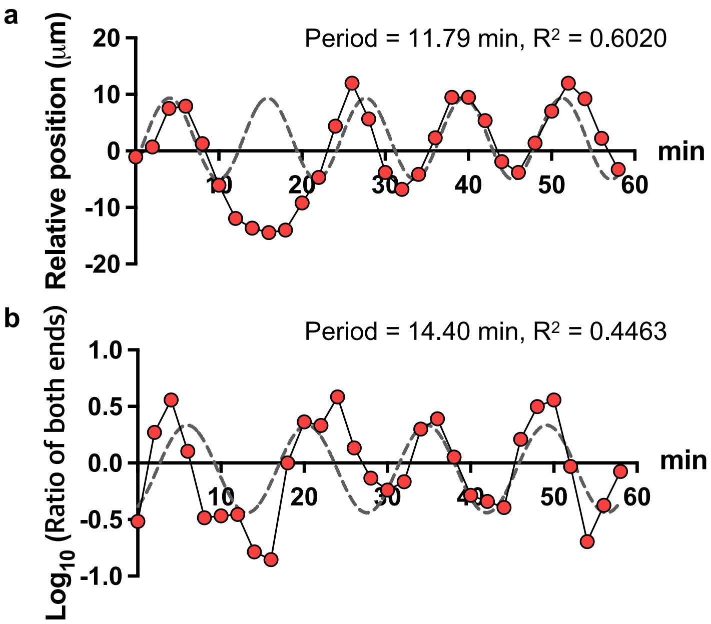


**Figure S6.** Dynamic features observed in the nucleus (a) and both end circumstance (b) in the oscillating microglia, corresponding to those of graphs in **Figures 2e** and **3c**, respectively. In (a, b), each measured data point (red circles) was fitted by dotted lines, respectively. Fitted values of the oscillating period in nuclear translocation (a; refer to **Figure 2e**) and the graphs (b; refer to **Figure 3c**) obtained from oscillating cells. The measured values and fitting curves were represented by red circles and gray-dotted lines, respectively.

### FIGURE S7

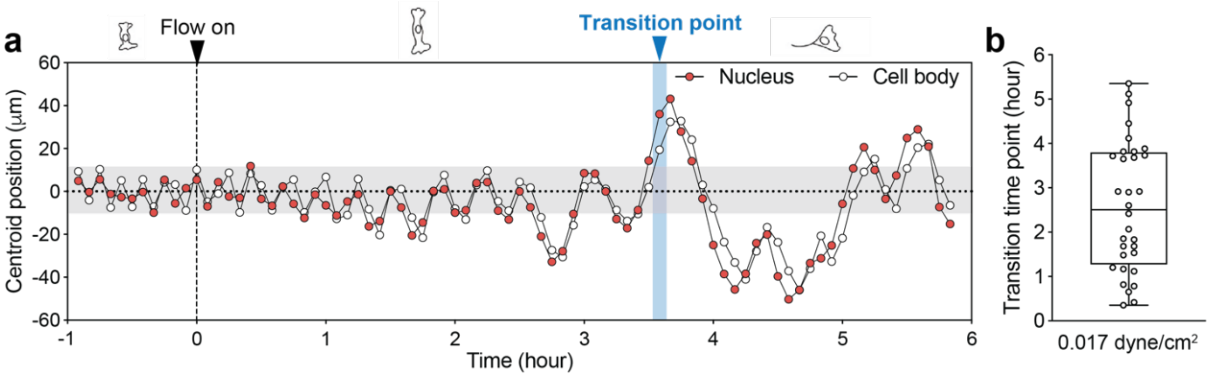


**Figure S7.** The transition from oscillatory to migratory microglia under the shear flow applied at 0.017 dyne/cm$^2$. (a) One representative example of shear-induced phenotypic transition. The centroid position of microglia was tracked at the time interval of 5 min for 6 hours. Here, the shear flow turned on at $t = 0$ hours, and the transition time point of the oscillatory microglia to migratory was indicated by an arrowhead at $t \sim$ 3.7 hours. (b) Scatter plot showing the distribution of transition time points observed for a total of 32 microglia.

### FIGURE S8

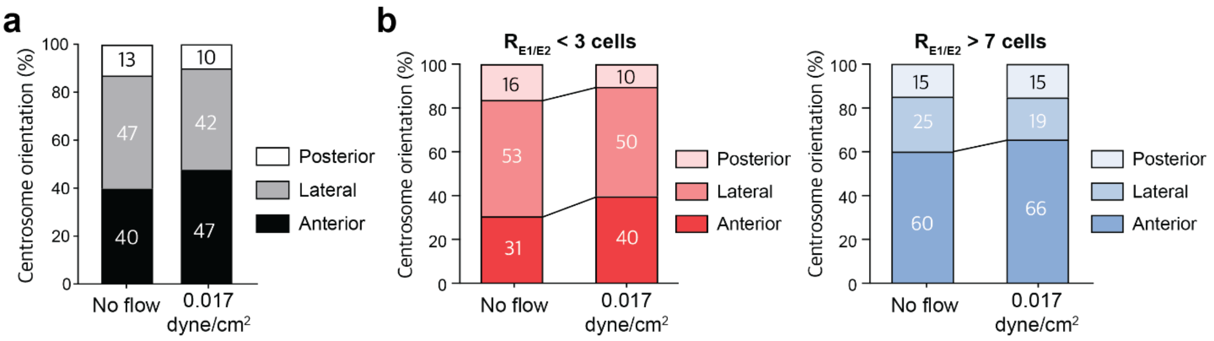


**Figure S8.** The orientation frequencies of the centrosome in microglia in the absence and presence of shear flow were analyzed from the immunostaining images of F-actin, g-tubulin, and DAPI. (a) The percentage of microglia having centrosomes in the anterior, lateral, and posterior. (b) The orientation frequencies of centrosomes in oscillating cells (polarity ratio < 3) and migrating cells (polarity ratio > 7).

**TABLE S1.** Primer sequences for gene analysis.

| Gene | Forward | Reverse |
|---|---|---|
| **GAPDH** | GGCAAGTTCAATGGCACAGT | TGGTGAAGACGCCAGTAGACTC |
| **IL-1β** | GCCCATCCTCTGTGACTCAT | AGGCCACAGGTATTTTGTCG |
| **TNF-α** | CGTCAGCCGATTTGCTATCT | CGGACTCCGCAAAGTCTAAG |
| **TGF-β1** | AAGAAGTCACCCGCGTGCTA | TGTGTGATGTCTTTGGTTTTGTCA |

**TABLE S2.** Fold change calculation. ΔCt = Ct value of target gene – Ct value of Gapdh; Fold change = $2^{-\Delta Ct}$/mean of $2^{-\Delta Ct}$ control. N = 5 for IL-1β and TNF-α groups, N = 4 for TGF-β1 and TREM2 groups.

| | | Control | | | | | Shear | | | | |
|---|---|---|---|---|---|---|---|---|---|---|---|
| **Genes** | | **Gapdh** | **IL-1β** | **TNF-α** | **TGF-β1** | **TREM2** | **Gapdh** | **IL-1β** | **TNF-α** | **TGF-β1** | **TREM2** |
| **Ct value** | 1 | 18.690 | 27.190 | 23.380 | 20.870 | 19.230 | 21.200 | 23.200 | 21.790 | 23.210 | 23.210 |
| | 2 | 19.070 | 27.360 | 25.430 | 21.690 | 19.430 | 18.880 | 18.740 | 18.380 | 21.360 | 21.600 |
| | 3 | 21.015 | 25.250 | 25.650 | 22.310 | 19.385 | 20.995 | 18.505 | 20.515 | 22.835 | 22.465 |
| | 4 | 20.120 | 23.520 | 24.320 | 21.780 | 19.250 | 19.885 | 18.035 | 19.200 | 21.935 | 22.595 |
| | 5 | 21.628 | 30.318 | 29.933 | n.a | n.a | 21.188 | 18.763 | 21.980 | n.a | n.a |
| **ΔCt** | 1 | | 8.500 | 4.690 | 2.180 | 0.540 | | 2.000 | 0.590 | 2.010 | 2.010 |
| | 2 | | 8.290 | 6.360 | 2.620 | 0.360 | | -0.140 | -0.500 | 2.480 | 2.720 |
| | 3 | | 4.235 | 4.635 | 1.295 | -1.630 | | -2.490 | -0.480 | 1.840 | 1.470 |
| | 4 | | 3.400 | 4.200 | 1.660 | -0.870 | | -1.850 | -0.685 | 2.050 | 2.710 |
| | 5 | | 8.690 | 8.305 | n.a | n.a | | -2.425 | 0.793 | n.a | n.a |
| **$2^{-\Delta Ct}$** | 1 | | 0.003 | 0.039 | 0.221 | 0.688 | | 0.250 | 0.664 | 0.248 | 0.248 |
| | 2 | | 0.003 | 0.012 | 0.163 | 0.779 | | 1.102 | 1.414 | 0.179 | 0.152 |
| | 3 | | 0.053 | 0.040 | 0.408 | 3.095 | | 5.618 | 1.395 | 0.279 | 0.361 |
| | 4 | | 0.095 | 0.054 | 0.316 | 1.828 | | 3.605 | 1.608 | 0.242 | 0.153 |
| | 5 | | 0.002 | 0.003 | n.a | n.a | | 5.370 | 0.577 | n.a | n.a |
| **mean** | | | 0.031 | 0.030 | 0.277 | 1.597 | | | | | |
| **Fold change** | 1 | | 0.088 | 1.302 | 0.797 | 0.431 | | 8.002 | 22.333 | 0.897 | 0.155 |
| | 2 | | 0.102 | 0.409 | 0.588 | 0.488 | | 35.269 | 47.542 | 0.648 | 0.095 |
| | 3 | | 1.700 | 1.353 | 1.472 | 1.938 | | 179.808 | 46.888 | 1.009 | 0.226 |
| | 4 | | 3.032 | 1.829 | 1.143 | 1.144 | | 115.385 | 54.047 | 0.872 | 0.096 |
| | 5 | | 0.078 | 0.106 | n.a | n.a | | 171.887 | 19.409 | n.a | n.a |
| **mean** | | | 1.000 | 1.000 | 1.000 | 1.000 | | 102.070 | 38.044 | 0.856 | 0.143 |
| **SEM** | | | 0.596 | 0.320 | 0.195 | 0.352 | | 34.933 | 7.137 | 0.076 | 0.031 |
